# Trait and state patterns of basolateral amygdala connectivity at rest are related to endogenous testosterone and aggression in healthy young women

**DOI:** 10.1101/248930

**Authors:** Macià Buades-Rotger, Christin Engelke, Ulrike M. Krämer

## Abstract

The steroid hormone testosterone (T) has been suggested to influence reactive aggression upon its action on the basolateral amygdala (BLA), a key brain region for threat detection. However, it is unclear whether T modulates resting-state functional connectivity (rsFC) of the BLA, and whether this predicts subsequent aggressive behavior. Aggressive interactions themselves, which often induce changes in T concentrations, could further alter BLA rsFC, but this too remains untested. Here we investigated the effect of endogenous T on rsFC of the BLA at baseline as well as after an aggressive encounter, and whether this related to behavioral aggression in healthy young women (n=39). Pre-scan T was negatively correlated with basal rsFC between BLA and left superior temporal gyrus (STG), which in turn predicted increased aggression. BLA-STG coupling at rest might thus underlie hostile readiness in low-T women. In addition, coupling between the BLA and the right superior parietal lobule (SPL), a brain region involved in higher-order perceptual processes, was reduced in aggressive participants. On the other hand, post-task increases in rsFC between BLA and medial orbitofrontal cortex (mOFC) were linked to reduced aggression, consistent with the established notion that the mOFC regulates amygdala activity in order to curb aggressive impulses. Finally, competition-induced changes in T were associated with increased coupling between the BLA and the right lateral OFC, but this effect was unrelated to aggression. We thus identified connectivity patterns that prospectively predict aggression in women, and showed how aggressive interactions in turn impact these neural systems.

## 1. Introduction

Human aggression has been often associated with enhanced responsivity to threat signals in the amygdala (Gospic et al. 2011; McCloskey et al. 2016; Buades-Rotger et al. 2016) as well as with lower orbitofrontal cortex (OFC) recruitment (Beyer et al. 2015; Mehta and Beer 2009). Concordantly, reduced coupling between these brain regions is thought to reflect deficient control of aggressive impulses (McCloskey et al. 2016; Coccaro et al. 2007; White et al. 2015; Wahlund and Kristiansson 2009). A relevant neuromodulator of this circuitry is the steroid hormone testosterone (T). T has been traditionally related to dominance and aggression, although some evidence indicates that this hormone can increase prosocial behavior in humans if it benefits one’s status - an effect that might be more pronounced in women (Eisenegger et al. 2011; Terburg and van Honk 2013). Regardless of its net effect on social behavior, T seems to mitigate fear responses (Terburg et al. 2016; Enter et al. 2016), at least in part by reducing threat-related coupling between the amygdala and the OFC (Bos et al. 2012; Spielberg et al. 2015). Previously, we observed that amygdala reactivity to angry faces was associated with lower T and increased aggression in the course of a competitive interaction in healthy young women (Buades-Rotger et al. 2016). This effect was maximal in the basolateral amygdala (BLA), a subregion involved in threat perception (de Gelder et al. 2014), and concurred with decreased BLA-OFC connectivity during trials in which the opponent displayed an angry face. In that study, as in most recent neuroscientific investigations on reactive aggression (Fanning et al. 2017), we focused on acute brain responses to threat signals. However, there might be relatively stable, trait-like neurofunctional characteristics that predispose individuals to react aggressively (Davidson et al. 2000; Fulwiler et al. 2012).

Such neural markers might be characterized by means of resting-state functional connectivity (rsFC), i.e. temporal correlations between remote brain regions at rest. Altered rsFC has been observed in different psychiatric conditions such as attention deficit hyperactivity disorder (ADHD; Castellanos and Aoki 2016), depression (Zhou et al. 2017), or autism (Hull et al. 2017), and normal variation in rsFC has been associated with personality characteristics such as neuroticism or extraversion (Aghajani et al. 2014). Importantly, dynamics of resting-state networks can also predict subsequent task performance (Sadaghiani et al. 2015; Fox et al. 2007), suggesting that rsFC not only captures stable differences between individuals, but also transient brain states that have an impact on behavior. Furthermore, there is substantial but not complete overlap between evoked functional networks across tasks and temporally coherent networks at rest (Cole et al. 2014; Mennes et al. 2013). This indicates that there is indeed an intrinsic neurofunctional architecture that can be nonetheless modulated by external stimulation. It is hence pertinent to investigate how task-evoked and resting-state networks relate to each other (Cole et al. 2016), and how they thereby influence behavior and cognition.

In the context of aggression, amygdala-OFC decoupling at rest has been observed in persons diagnosed with Borderline Personality Disorder (BPD; New et al. 2007; Baczkowski et al. 2016) and in children with ADHD (Ho et al. 2015), patient groups that often display elevated aggressiveness (Prada et al. 2014). Lower amygdala-OFC rsFC has also been associated with self-reported aggression in male schizophrenia patients (Hoptman et al. 2010), with psychopathy (Motzkin et al. 2011), and with trait anger in healthy participants (Fulwiler et al. 2012). T could play an additional role in this neural circuitry and in turn in aggression by modulating not only threat reactivity of the amygdala, but also its rsFC (Savic et al. 2017). For instance, anabolic androgen users, who are often aggression-prone (Pope et al. 2014), show reduced rsFC between the amygdala and the so-called default mode network (DMN), a set of regions that includes the medial OFC, the precuneus, the superior temporal gyrus (STG), and other areas involved in social cognition (Westlye et al. 2017). Comparably, endogenous T has been associated with heightened alcohol use - an established risk factor for aggression (Tremblay et al. 2008) - in subjects with low amygdala-OFC connectivity at rest (Peters et al. 2015).

The reviewed findings indicate that basal patterns of rsFC are modulated by T and can predict aggression. Nonetheless, most of these studies rely on self-reports, which are strongly affected by social desirability (Vigil-Colet et al. 2012). It remains to be seen whether comparable results can be obtained with behavioral measures of aggression, which should be less affected by this confound as well as by linguistic or interpretational biases. Indeed, behavioral aggression measures have been shown to have high internal and external validity (Anderson and Bushman 1997; Giancola and Parrott 2008), and to better capture links between T and aggression than self-reports (Archer et al. 2005).

It is also unclear whether and to which extent rsFC patterns are in turn modulated by aggressive encounters themselves. In this vein, some researchers have probed for changes in resting-state networks after stressful social situations. These studies have shown that social exclusion increases connectivity between the DMN and brain areas involved in threat detection and conflict monitoring (Clemens et al. 2017), and that evaluative stress changes thalamic-cortical and parietal-temporal coupling (Maron-Katz et al. 2016). In such investigations, however, subjects were passively exposed to a menacing situation, and so results cannot be readily extrapolated to interactive, reciprocal aggressive interactions. Moreover, there is disagreement regarding whether aggression is better predicted by trait or state estimates of T (Carré and Olmstead 2015). For instance, a recent study found that aggressive behavior was linked with lower basal T but higher T reactivity, and that the strength of these relationships depended on self-construal (Welker et al. 2017). Investigating the possible connection between state changes in T and aggression-related rsFC patterns can help to elucidate this issue.

Here, we analyzed resting-state data obtained in a previous study (Buades-Rotger et al. 2016) in order to answer two main questions. First, we attempted to ascertain whether basal rsFC of the amygdala can predict subsequent aggression as measured in a realistic task, and whether this was related to baseline T levels in healthy young women. Second, we inquired as to whether an aggressive interaction itself can induce changes in amygdala rsFC, and whether these putative alterations are likewise related to aggression and to state changes in T concentrations.

## 2. Methods

### 2.1 Participants

Participants were the same 39 healthy young women (mean age = 23.22, SD = 3.2) from our previously published study (Buades-Rotger et al. 2016). Exclusion criteria were history of neurologic, psychiatric, or endocrine disease and intake of oral contraceptives. All subjects were right handed and fluent German speakers. Informed consent was obtained from all participants, who were also compensated for participation. The study was performed in accordance with the Declaration of Helsinki and had been approved by the University of Lübeck ethics committee.

### 2.2 Procedure

On the first measurement day subjects provided informed consent and underwent an anatomical scan in order to accustom them to the scanner environment. Participants also provided ambulatory saliva samples during a regular weekday. The second measurement day was always scheduled during the early follicular phase and in the afternoon (i.e. between noon and 6 pm) in order to limit daily (Keevil et al. 2013) and monthly (Bui et al. 2013) variability in hormonal concentrations. On this second measurement occasion, participants provided a blood and a saliva sample before scanning. We acquired two 6-minute, eyes-closed resting state runs *before* and *after* the aggression task, which was named Social Threat Aggression Paradigm (STAP). In the STAP, participants face a same-sex opponent who displays an angry or a neutral face at the beginning of each trial and provokes them with progressively louder sound blasts. Participants must choose the intensity of a sound blast in a scale of 1 to 8 to be directed at the opponent in case of victory. Actually, the game is preprogrammed and the angry and neutral videos are prerecorded. After scanning, participants provided a second saliva sample and fulfilled a series of personality questionnaires not analyzed here (see Buades-Rotger et al. 2016). Finally, subjects were probed for suspicion, debriefed, and compensated.

### 2.3 Hormone data acquisition

Participants provided saliva samples during a regular weekday as well as before and after scanning using the passive drool method (Granger et al. 2007). We asked them to fill at least 3 mL of each tube (4 mL Cryovials from Salimetrics ®), and to refrain from: a) eating or teeth brushing within 1 hour prior to collection, b) using salivation stimulants such as chewing gum, c) drinking alcohol within 12 hours prior to collection, and d) going to the dentist within 48 h prior to collection. Samples were stored at ‐20°C and shipped in dry ice-filled Styrofoam boxes to a reference laboratory at University Hospital of South Manchester (UK) for analysis. Free salivary T concentrations were estimated using a liquid chromatography tandem mass spectrometry (LC-MS/MS) method optimized to capture values in the low range (Keevil et al. 2013). As previously reported, mean intra‐ and inter-assay coefficients of variation for salivary T were 5.3% and 9%, with a lower limit of quantification of 5 pmol/L (Keevil et al. 2013). Serum T and salivary cortisol were also estimated, but we focused our present analyses on T at scan-time because it was the only hormonal parameter associated with aggression and BLA reactivity (Buades-Rotger et al. 2016). T concentrations were log-transformed to stabilize variance before analysis.

### 2.4 Neuroimaging data acquisition

Neuroimaging data was acquired using a Philips Ingenia 3T scanner equipped with a 32-channel head coil. The anatomical scan was performed with a standard T1-weighted EPI sequence (180 sagittal slices, TR=7.7, TE=3.5, FOV=240, matrix=240 x 240 mm, flip angle=8°, voxel size=1 mm isotropic). Resting-state data was acquired with a T2*-weighted gradient EPI sequence sensitive to the blood-oxygen level-dependent (BOLD) signal (37 axial slices per volume, TR=2 s; TE=30 ms; FOV=192 mm, matrix=64 x 64 mm; flip angle=90°; voxel size=3 mm isotropic). We acquired 178 volumes plus 5 dummy scans before and after the task.

### 2.5 Data analysis

#### 2.5.1. Behavioral data analysis

Behavioral results have already been reported (Buades-Rotger et al. 2016), but we will briefly explain how aggression data was analyzed for ease of interpretation. We used repeated measures analyses of variance (ANOVA) to test whether participants became more aggressive over the three runs of the task and in angry versus neutral trials, and whether they selected higher punishments after high (>4) relative to low (≤4) punishment selections by the opponent. Since excessive analytic flexibility can be problematic when extracting behavioral indexes from the competitive reaction time task (Elson et al. 2014), we took average punishment selections over the course of the task for each participant as a simple summary measure of reactive aggression for further analyses, as in our previous studies (Buades-Rotger et al. 2016; Beyer et al. 2015).

#### 2.5.2. Neuroimaging data preprocessing

Initial preprocessing of the data was performed with Statistical Parametric Mapping 12 (SPM12, Wellcome Department of Imaging Neuroscience, University College London, London, UK) and comprised slice-timing correction to the middle slice, realignment to the first volume, segmentation of the anatomical image, coregistration of mean functional and anatomical images, normalization to Montreal Neurological Institute (MNI) space and smoothing with an 8 mm full width at half-maximum (FWHM) Gaussian kernel. White matter (WM) and cerebrospinal fluid (CSF) masks obtained from segmentation were also normalized to MNI space. Additional denoising and subsequent rsFC analyses were performed with CONN v17 (Whitfield-Gabrieli and Nieto-Castanon 2012). Denoising involved regressing out signal from individual white matter and CSF masks as well as from movement parameters, linear detrending, despiking, and bandpass filtering (0.008-0.1 Hz) in order to isolate the low frequency range at which intrinsic connectivity operates (Zuo et al. 2010; Buckner et al. 2013).

#### 2.5.4. Neuroimaging data analyses

Both the BLA seed and the OFC region-of-interest (ROI) were informed by task activation. For ROI-to-ROI analyses, we used the same BLA and medial OFC (mOFC) masks that showed reduced reciprocal connectivity when the opponent displayed angry faces during the STAP (Buades-Rotger et al. 2016). The bilateral BLA mask covered all voxels activated in the decision phase, whereas the mOFC ROI was an 8-mm sphere around the group peak that was significantly more active for angry than for neutral faces. For whole-brain analyses, we created an anatomical mask covering the whole BLA using the Anatomy toolbox (Eickhoff et al. 2007). This seed was largely overlapping with the functional amygdala mask, but contained more voxels and hence provided increased power to detect effects. Using an anatomical mask also improves comparability with other rsFC studies on amygdala subregions (Aghajani et al. 2016; Qin et al. 2014; Roy et al. 2009).

Connectivity was computed as the Fisher-transformed correlation coefficient between the mean BOLD signal in the functional BLA mask and the mOFC for ROI-to-ROI analyses, or between the mean BOLD signal in the anatomical BLA mask and the rest of the brain for whole-brain analyses.

We first probed whether trait estimates of BLA rsFC were associated with basal T and aggression. For ROI analyses, we computed Pearson correlation coefficients between pre-task BLA-mOFC coupling, aggression, and pre-scan T across participants. For whole-brain analyses, we regressed pre-scan T values and aggression against pre-task BLA connectivity.

We then tested whether there were task-induced alterations in BLA rsFC, and whether these were related to aggression and state changes in T. At the ROI level, we compared BLA-mOFC connectivity estimates before and after the task with a paired t-test, and correlated this difference with aggression and with the post-minus prescan change in T. At the whole-brain level, we compared post vs. pre-task BLA rsFC with a paired t-test, and regressed aggression and the change in T against the post-minus pre-task difference in BLA rsFC.

Additionally, we explored whether regions showing T-modulated connectivity with the BLA in whole-brain analyses were associated with aggression. We thus extracted connectivity estimates from T-related clusters and inspected for correlations with aggression.

Whole-brain analyses were thresholded at p<.001 uncorrected at the voxel level with a family-wise error (FWE) correction of p<.05 at the cluster level, whereas ROI analyses were deemed significant at p<.05 (uncorrected).

### 3. Results

#### 3.1 Behavioral results

Behavioral results are described in detail in Buades-Rotger, et al. (2016), but we report the essential effects again here for completeness. The STAP was successful in provoking participants, eliciting higher aggression over runs (F_2,76_=21.93, p<.001; Run 1=3.5±0.12 [mean ± standard error], Run 3=4.3±0.14) and in angry compared to neutral trials (F_1, 38_=6.34, p=.016; Angry=4.13±0.16; Neutral=3.94±0.14). Furthermore, participants reacted to the opponent’s aggression (F_1, 38_=20.31, p<.001) by choosing stronger punishments after high (4.27±0.10) than after low (3.85±0.09) punishment selections. As in previous work (Beyer et al. 2015; Buades-Rotger et al. 2016), we extracted each participant’s mean punishment selections as our main aggression measure for further analyses.

### 3.2 Associations between basal BLA rsFC, T and aggression

BLA-mOFC coupling was unrelated to T (p=.916) and aggression (p=.240) at baseline. In whole-brain analyses, pre-scan T was associated with lower baseline coupling between the BLA and the left superior temporal gyrus (STG; peak MNI coordinates: ‐33, ‐28, 2; k=61; peak T=4.25; cluster-level pFWE=.016; Fig. 1A and 1B). BLA-STG connectivity strength was further associated with increased aggression (r=.37, p=.020; Fig. 1C), such that participants with high BLA-STG coupling had reduced T levels and were more aggressive. The STG cluster was in the inner section of the temporal operculum, around Brodmann area 41 extending into the posterior insular cortex. It was adjacent to, though slightly more medially located than the aggression-related left STG cluster identified in Buades-Rotger et al. (2016).

**Fig. 1.**
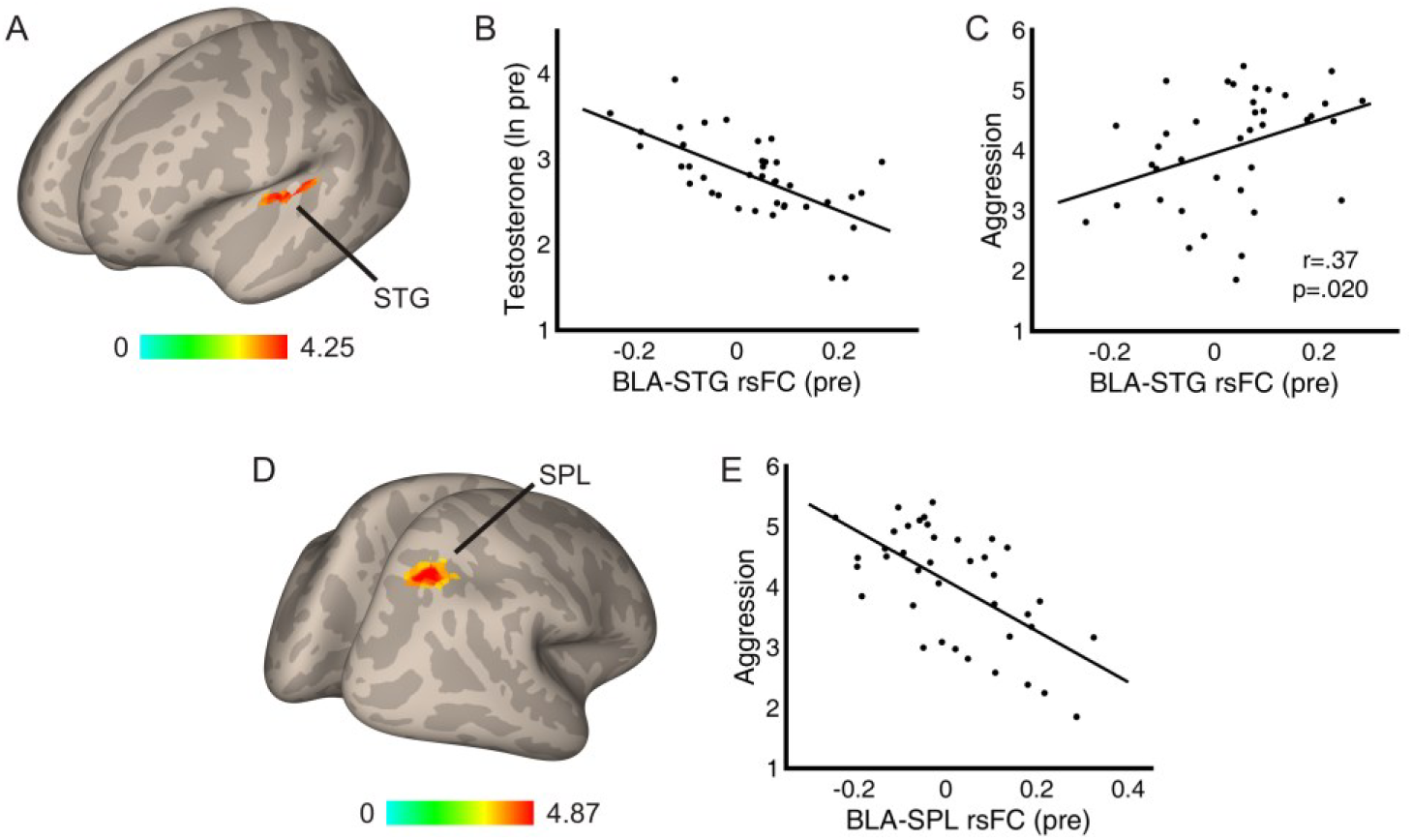
Trait resting-state functional connectivity (rsFC) of the basolateral amygdala (BLA) was associated with testosterone (T) and aggression. **A:** Left superior temporal gyrus (STG) cluster showing lower pre-task coupling with BLA as a function of basal T (p<.001, pFWE<.05 cluster-level corrected). **B:** scatterplot and least-squares fit line of the correlation between BLA-STG coupling and log-transformed T concentrations (r and p values not computed to avoid circular analysis). **C**: scatterplot and least-squares fit line of the negative correlation between BLA-STG coupling and aggression. Participants with higher basal BLA-STG coupling had lower pre-scan T and were more aggressive. **D:** cluster in the right superior parietal lobule (SPL) showing reduced pre-task rsFC with the BLA as a function of aggression (p<.001, pFWE<.05 cluster-level corrected). **E:** scatterplot and least-squares fit line of the negative correlation between pre-task BLA-SPL rsFC and aggression (r and p values not computed to avoid circular analysis). Participants with stronger BLA-SPL rsFC were less aggressive.

Basal connectivity between the BLA and the superior parietal lobule (SPL) was associated with lower aggression (peak MNI coordinates: 27, ‐61, 47; k=51; peak T=4.87; cluster-level pFWE=.035; Fig. 1D and 1E). The local maximum was located in Brodmann area 7, though the cluster covered the ventral-posterior aspect of the superior parietal lobule extending to the intraparietal sulcus.

### 3.3 Associations between state changes in BLA rsFC, T and aggression

There was no significant change in BLA-mOFC coupling after compared to before the task (p=.664), and difference values were not associated with the change in T (p=.819). However, the post-minus pre-task difference in rsFC between these regions was negatively associated with aggression (r=-.36, p=.023; Fig. 2A and 2B). Participants with higher increases in BLA-mOFC connectivity were less aggressive. The relationship was stronger for post‐ (r=-.30, p=.058) than for pre-task coupling (r=.19, p=.240), implying that the effect was mostly driven by a task-induced increase in connectivity rather than by high baseline values.

**Fig. 2.**
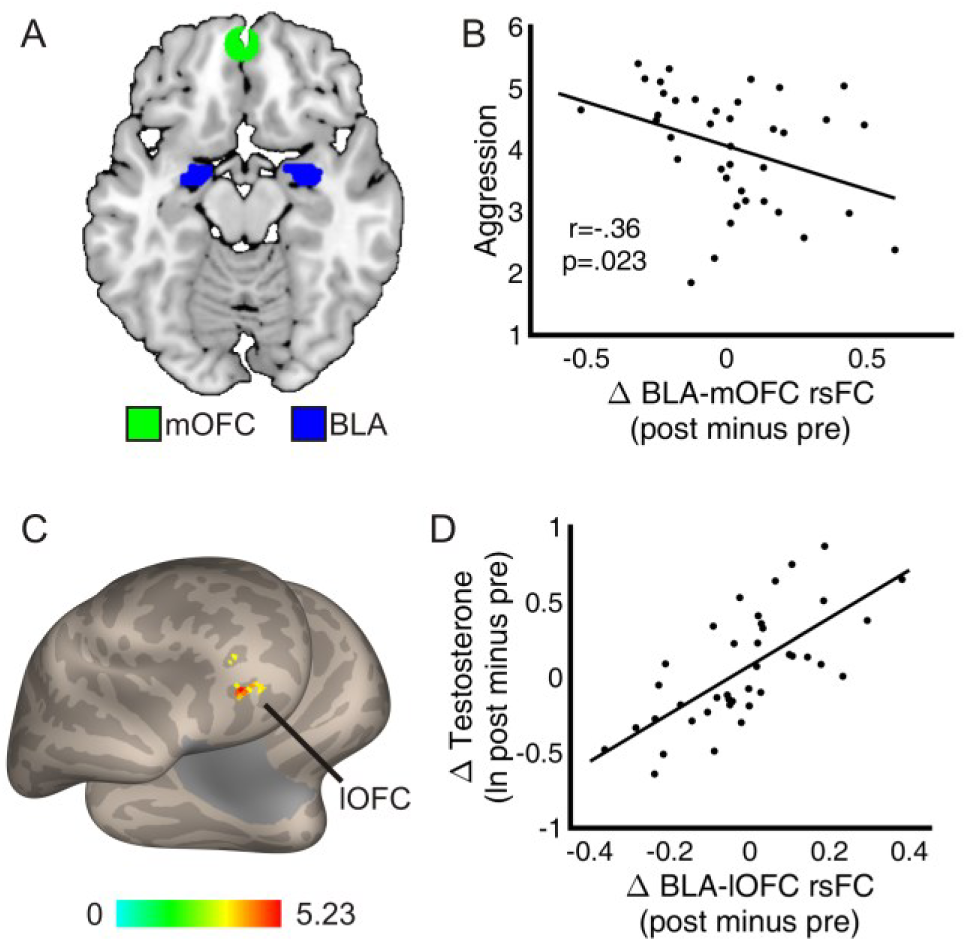
State changes in resting-state functional connectivity (rsFC) of the basolateral amygdala (BLA) were related to testosterone (T) and aggression. **A:** functionally defined seeds in BLA (blue) and medial orbitofrontal cortex (mOFC; green). **B:** scatterplot and least-squares fit line of the negative correlation between task-induced change in BLA-mOFC rsFC and aggression. Participants with higher post-relative to pre-task BLA-OFC connectivity were less aggressive. **C:** cluster in the right lateral orbitofrontal cortex (lOFC) showing reduced coupling with the BLA as a function of the post-minus pre-scan change in T (p<.001, pFWE<.05 cluster-level corrected). **D:** scatterplot and least-squares fit line of the positive correlation between the post-minus pre-task difference in BLA-lOFC coupling and the log-transformed change in T concentrations (r and p values not computed to avoid circular analysis). Participants with higher post-relative to pre-task BLA-lOFC rsFC showed stronger increases in T.

There were no significant differences between post‐ and pre-task BLA connectivity in whole-brain analyses. Nonetheless, the change in T concentrations was correlated with higher post-minus pre-task coupling between the BLA and the right lateral OFC (lOFC; peak MNI coordinates: 36, 38, ‐7; k=44; peak T=5.51; cluster-level pFWE=.048; Fig. 2C and 2D). That is, participants with a stronger increase in T showed higher BLA-lOFC coupling after the task. However, this effect was unrelated to aggression (p=.514). The lOFC cluster was located between Brodmann areas 11 and 47.

## 4. Discussion

We investigated whether intrinsic connectivity of the BLA was associated with trait and state estimates of T, and whether this could predict aggressive behavior in a sample of healthy young women. We found that pre-scan T was negatively correlated with BLA-STG connectivity, which in turn predicted higher aggression during the task. Basal coupling between the BLA and the SPL predicted lower aggression. On the other hand, task-induced changes in rsFC between the BLA and the mOFC were associated with reduced aggression, whereas T changes were linked to higher BLA-lOFC coupling. BLA-STG and BLA-SPL rsFC thus seem to constitute stable features that distinguish aggressive from non-aggressive individuals, whereas BLA-OFC connectivity at rest might be more volatile and dependent on the organism’s reactions to threat.

### 4.1 BLA rsFC is related to baseline T and predicts subsequent aggression

Basal rsFC between the BLA and the left STG was inversely correlated with T, and positively associated with aggressive behavior. This dovetails with our previously published task results showing that STG activation in response to angry faces mediates the relationship between BLA reactivity and aggression (Buades-Rotger et al. 2016). The present data suggests that BLA-STG coupling is not only important for momentary threat assessment, but might also underlie hostile readiness. In agreement with this notion, rsFC between the BLA and a frontotemporal network including the STG has been related to self-reported impulsivity in children with conduct disorder (Aghajani et al. 2016). Furthermore, the amygdala and the STG are co-recruited during the perception of angry vocalizations (Mothes-Lasch et al. 2012; Frühholz et al. 2015), of threatening facial expressions (Sussman et al. 2016), and of anger-infused body movements (Grèzes et al. 2013; Van den Stock et al. 2015). Although the localization of the current STG cluster (neighboring Heschl’s gyrus) is unusual, this medial aspect of the STG extending to the posterior insula (PI) has been shown to react preferentially to threatening relative to happy, neutral or sad stimuli independently of the sensory modality (i.e. fearful faces, fear-inducing music and screams; Aubé et al. 2015). In that study, the observed peak (MNI coordinates: ‐38, ‐26, 2) was no more than 5 mm away in any direction from the current one (MNI coordinates: ‐33, ‐28, 2), and, crucially, the BLA showed a highly similar pattern of activity as that of the STG. Note also that the aggression-related temporal clusters identified in our previous study were likewise in the medial section of the STG (Buades-Rotger et al. 2016). Taken together, this evidence indicates that the assumed hostile readiness observed in individuals with high BLA-STG coupling might be characterized by increased attunement towards multimodal threat signals.

Crucially, BLA-STG coupling was reduced in women with higher T levels, who were also less aggressive and showed lower amygdala reactivity to angry faces during the STAP (Buades-Rotger et al. 2016). The STG seems to be particularly sensitive to T, as suggested by studies in female-to-male transsexuals which reported lower gray matter volume in this brain area after long-term T administration (A. Hahn et al. 2016; Burke et al. 2017). In line with our results, long-term users of anabolic androgens show generally blunted amygdala rsFC with a widespread network that includes the medial prefrontal cortex and the STG relative to matched non-users (Kaufman et al. 2015; Westlye et al. 2017). This pattern also fits studies suggesting that T reduces threat sensitivity (Terburg et al. 2016; Enter et al. 2016) and decreases connectivity within emotion-processing networks (Bos et al. 2016; van Wingen et al. 2010). All in all, as reasoned above, it seems that the basal interplay between BLA and STG might reflect a lower threshold for cross-modal threat perception and might hence facilitate corresponding aggressive responses in low-T women.

In addition, pre-task coupling between the BLA and the SPL was inversely correlated with aggression. The SPL subregion identified here corresponds to clusters 3 and 4 identified in Wang et al. (2015), which are roughly equivalent to cytoarchitectonic areas hIP3 and 7A (Scheperjans et al. 2008). Across studies, these subregions are recruited when performing complex cognitive tasks, most notably those involving spatial attention allocation, reasoning, and working memory (Wang et al. 2015). Although replication is warranted, we speculate that a higher basal BLA-SPL coupling could be indicative of relatively improved cognitive ability, which might contribute to successful emotion regulation. The finding that depressed patients (Chen et al. 2017) and individuals with high dispositional sadness (Deris et al. 2017) show reduced amygdala-SPL coupling at rest agrees with this formulation.

### 4.1 Changes in BLA-OFC rsFC induced by an aggressive interaction

Post-task increments in BLA-mOFC rsFC were related to lower aggression. As reviewed in the introduction, reduced rsFC between these two regions is generally associated with impaired impulse control and antisocial behavior (Peters et al. 2016; Hoptman et al. 2010; Fulwiler et al. 2012; Motzkin et al. 2011). Our data is consistent with these results and fit the widely accepted view that the OFC regulates amygdala activity in order to curb aggressive behavior (Coccaro et al. 2011). The present study further adds that this mechanism is not only manifest in basal connectivity patterns or in transient threat-related coupling, but that positive shifts in BLA-mOFC rsFC induced by a challenging social encounter might also support the control of aggressive impulses. Complementing our results, a recent study found that healthy participants show increased rsFC between the amygdala and the OFC after an emotion regulation task, whereas BPD patients failed to show such an increase (Baczkowski et al. 2016).

It is noteworthy that we did not observe any relationship between amygdala-mOFC rsFC and endogenous T. We did however find a positive relationship between state changes in T and increases in coupling between the BLA and the lOFC. This might be due to the relatively higher density of androgen receptors in the lateral relative to the medial wall of the prefrontal cortex (Low et al. 2017). In any case, our results contrast with previous studies showing threat-related decreases in amygdala-OFC coupling concurrent with increases in T, both acutely (van Wingen et al. 2010) and longitudinally (Spielberg et al. 2015). The observed increase in BLA-lOFC coupling at rest might thus conceivably arise as a post-encounter recovery mechanism in participants with stronger changes in T, who were presumably more competitive (A. C. Hahn et al. 2016; Casto et al. 2017) and might have been therefore more engaged in the task. Indirectly supporting this notion, increased task-related coupling between the amygdala and the lOFC has been related to reduced negative affect in healthy participants (Madsen et al. 2016; Banks et al. 2007), to diminished impulsivity in psychopaths (Yoder et al. 2015), and to lower state anger in BPD patients (Herpertz et al. 2017), a clinical group with heightened endogenous T levels (Rausch et al. 2015). Nevertheless, given that BLA-lOFC rsFC was not related to aggression, it is difficult to interpret the possible cognitive or behavioral significance of this functional link. Note also that T concentrations did not change significantly after the task and the post-minus pre-scan difference in T was unrelated to aggression (Buades-Rotger et al. 2016). Furthermore, the effects of competition on T levels are relatively unstable in women (Carré and Olmstead 2015), which calls for additional caution when interpreting this result.

### 4.3 Limitations

Certain limitations of the present study should be taken into account. Some are inherent to rsFC, such as the non-directionality of the observed functional links and the unconstrained cognitive activity during scanning at rest (Buckner et al. 2013). Other shortcomings are specific for this study. The relatively small sample size was lower than what has become standard in recent rsFC studies. However, our dataset is unique in that we employed a realistic behavioral measure of aggression, which, despite having limitations of its own (Ritter and Eslea 2005), is more ecologically valid and less affected by psychometric biases than questionnaires (Anderson and Bushman 1997). Using such a task also allowed us to investigate how an aggressive interaction itself alters rsFC of the amygdala, which, to our knowledge, had not been investigated before. Another limitation is that we selected our ROIs on the basis of task results, which might be less optimal than individually delineated ROIs (Sohn et al. 2015). This might have curtailed our ability to detect effects, but does in fact support the robustness of the effects that we did observe.

### 4.4 Conclusion

In summary, we found that BLA-STG rsFC might underlie hostile readiness in anticipation of an aggressive encounter, and we showed that T might buffer aggression in women by weakening this link. Coupling between the BLA and the SPL might oppositely constitute a regulatory mechanism. Finally, we observed a dissociation between lateral and medial OFC, such that stronger post-task BLA-lOFC was related to competition-induced changes in T, whereas increases in BLA-mOFC coupling were associated with reduced aggression. All in all, our study identifies how aggressive interactions plastically change amygdala connectivity, and pinpoint specific neural mechanisms through which T can buffer aggression in women.

## Acknowledgements

This study was funded by the German Science Foundation (grant number KR3691/5-1). We are grateful to Matthias Liebrand, Alice Nöh, Christian Erdmann, and Susanne Schellbach for their help with the data acquisition. The authors report no conflicts of interest.

## References

Aghajani, M., Colins, O. F., Klapwijk, E. T., Veer, I. M., Andershed, H., Popma, A., et al. (2016). Dissociable relations between amygdala subregional networks and psychopathy trait dimensions in conduct-disordered juvenile offenders. Humanbrain mapping, 37(11), 4017-4033, doi:10.1002/hbm.23292.

Aghajani, M., Veer, I. M., van Tol, M.-J., Aleman, A., van Buchem, M. A., Veltman, D. J., et al. (2014). Neuroticism and extraversion are associated with amygdala resting-state functional connectivity. Cognitive, Affective, & Behavioral Neuroscience, 14(2), 836-848, doi:10.3758/s13415-013-0224-0.

Anderson, C. A., & Bushman, B. J. (1997). External validity of “trivial” experiments: The case of laboratory aggression. Review of General Psychology, 1(1), 19-41.

Archer, J., Graham-Kevan, N., & Davies, M. (2005). Testosterone and aggression: A reanalysis of Book, Starzyk, and Quinsey's (2001) study. Aggression and Violent Behavior, 10(2), 241-261, doi:http://dx.doi.org/10.1016/j.avb.2004.01.001.

Aubé, W., Angulo-Perkins, A., Peretz, I., Concha, L., & Armony, J. L. (2015). Fear across the senses: brain responses to music, vocalizations and facial expressions. Social Cognitive and Affective Neuroscience, 10(3), 399-407, doi:10.1093/scan/nsu067.

Baczkowski, B. M., van Zutphen, L., Siep, N., Jacob, G. A., Domes, G., Maier, S., et al. (2016). Deficient amygdala–prefrontal intrinsic connectivity after effortful emotion regulation in borderline personality disorder. European Archives of Psychiatry and Clinical Neuroscience, 1-15, doi:10.1007/s00406-016-0760-z.

Banks, S. J., Eddy, K. T., Angstadt, M., Nathan, P. J., & Phan, K. L. (2007). Amygdala-frontal connectivity during emotion regulation. Social Cognitive and Affective Neuroscience, 2(4), 303-312, doi:10.1093/scan/nsm029.

Beyer, F., Münte, T. F., Göttlich, M., & Krämer, U. M. (2015). Orbitofrontal Cortex Reactivity to Angry Facial Expression in a Social Interaction Correlates with Aggressive Behavior. Cerebral Cortex, 25(9), 3057-3063, doi:10.1093/cercor/bhu101.

Bos, P. A., Hofman, D., Hermans, E. J., Montoya, E. R., Baron-Cohen, S., & van Honk, J. (2016). Testosterone reduces functional connectivity during the ‘Reading the Mind in the Eyes’ Test. Psychoneuroendocrinology, 68, 194-201, doi:http://dx.doi.org/10.1016/j.psyneuen.2016.03.006.

Bos, P. A., Panksepp, J., Bluthé, R.-M., & Honk, J. v. (2012). Acute effects of steroid hormones and neuropeptides on human social-emotional behavior: A review of single administration studies. Frontiers in Neuroendocrinology, 33(1), 17-35, doi:10.1016/j.yfrne.2011.01.002.

Buades-Rotger, M., Engelke, C., Beyer, F., Keevil, B. G., Brabant, G., & Krämer, U. M. (2016). Endogenous testosterone is associated with lower amygdala reactivity to angry faces and reduced aggressive behavior in healthy young women. [Article]. Scientific Reports, 6, 38538, doi:10.1038/srep38538.

Buckner, R. L., Krienen, F. M., & Yeo, B. T. T. (2013). Opportunities and limitations of intrinsic functional connectivity MRI. [Review]. Nat Neurosci, 16(7), 832-837, doi:10.1038/nn.3423.

Bui, H. N., Sluss, P. M., Blincko, S., Knol, D. L., Blankenstein, M. A., & Heijboer, A. C. (2013). Dynamics of serum testosterone during the menstrual cycle evaluated by daily measurements with an ID-LC-MS/MS method and a 2nd generation automated immunoassay. Steroids, 78(1), 96-101, doi:10.1016/j.steroids.2012.10.010.

Burke, S. M., Manzouri, A. H., Dhejne, C., Bergstrom, K., Arver, S., Feusner, J. D., et al. (2017). Testosterone Effects on the Brain in Transgender Men. Cerebral Cortex, 1-15, doi:10.1093/cercor/bhx054.

Carré, J. M., & Olmstead, N. A. (2015). Social neuroendocrinology of human aggression: Examining the role of competition-induced testosterone dynamics. Neuroscience, 286, 171-186, doi:10.1016/j.neuroscience.2014.11.029.

Castellanos, F. X., & Aoki, Y. (2016). Intrinsic Functional Connectivity in Attention-Deficit/Hyperactivity Disorder: A Science in Development. Biological Psychiatry: Cognitive Neuroscience and Neuroimaging, 1(3), 253-261, doi:http://dx.doi.org/10.1016/j.bpsc.2016.03.004.

Casto, K. V., Rivell, A., & Edwards, D. A. (2017). Competition-related testosterone, cortisol, and perceived personal success in recreational women athletes. Hormones and Behavior, doi:https://doi.org/10.1016/j.yhbeh.2017.05.006.

Chen, H. J., Wang, Y. F., Qi, R., Schoepf, U. J., Varga-Szemes, A., Ball, B. D., et al. (2017). Altered Amygdala Resting-State Functional Connectivity in Maintenance Hemodialysis End-Stage Renal Disease Patients with Depressive Mood. Molecular Neurobiology, 54(3), 2223-2233, doi:10.1007/s12035-016-9811-8.

Clemens, B., Wagels, L., Bauchmüller, M., Bergs, R., Habel, U., & Kohn, N. (2017). Alerted default mode: functional connectivity changes in the aftermath of social stress. Scientific Reports, 7, 40180, doi:10.1038/srep40180.

Coccaro, E. F., McCloskey, M. S., Fitzgerald, D. A., & Phan, K. L. (2007). Amygdala and Orbitofrontal Reactivity to Social Threat in Individuals with Impulsive Aggression. Biological Psychiatry, 62(2), 168-178, doi:http://dx.doi.org/10.1016/j.biopsych.2006.08.024.

Coccaro, E. F., Sripada, C. S., Yanowitch, R. N., & Phan, K. L. (2011). Corticolimbic Function in Impulsive Aggressive Behavior. Biological Psychiatry, 69(12), 1153-1159, doi:10.1016/j.biopsych.2011.02.032.

Cole, Michael W., Bassett, Danielle S., Power, Jonathan D., Braver, Todd S., & Petersen, Steven E. (2014). Intrinsic and Task-Evoked Network Architectures of the Human Brain. Neuron, 83(1), 238-251, doi:http://dx.doi.org/10.1016/j.neuron.2014.05.014.

Cole, Michael W., Ito, T., Bassett, Danielle S., & Schultz, D. H. (2016). Activity flow over resting-state networks shapes cognitive task activations. Nature Neuroscience, 19(12), 1718-1726, doi:10.1038/nn.4406.

Davidson, R. J., Putnam, K. M., & Larson, C. L. (2000). Dysfunction in the Neural Circuitry of Emotion Regulation‐‐A Possible Prelude to Violence. Science, 289(5479), 591-594, doi:10.1126/science.289.5479.591.

de Gelder, B., Terburg, D., Morgan, B., Hortensius, R., Stein, D. J., & van Honk, J. (2014). The role of human basolateral amygdala in ambiguous social threat perception. Cortex, 52, 28-34, doi:http://dx.doi.org/10.1016/j.cortex.2013.12.010.

Deris, N., Montag, C., Reuter, M., Weber, B., & Markett, S. (2017). Functional connectivity in the resting brain as biological correlate of the Affective Neuroscience Personality Scales. NeuroImage, 147, 423-431, doi:https://doi.org/10.1016/j.neuroimage.2016.11.063.

Eickhoff, S. B., Paus, T., Caspers, S., Grosbras, M.-H., Evans, A. C., Zilles, K., et al. (2007). Assignment of functional activations to probabilistic cytoarchitectonic areas revisited. NeuroImage, 36(3), 511-521, doi:http://dx.doi.org/10.1016/j.neuroimage.2007.03.060.

Eisenegger, C., Haushofer, J., & Fehr, E. (2011). The role of testosterone in social interaction. Trends in Cognitive Sciences, 15(6), 263-271, doi:http://dx.doi.org/10.1016/j.tics.2011.04.008.

Elson, M., Mohseni, M. R., Breuer, J., Scharkow, M., & Quandt, T. (2014). Press CRTT to measure aggressive behavior: The unstandardized use of the competitive reaction time task in aggression research. Psychological Assessment, 26(2), 419-432, doi:10.1037/a0035569.

Enter, D., Terburg, D., Harrewijn, A., Spinhoven, P., & Roelofs, K. (2016). Single dose testosterone administration alleviates gaze avoidance in women with Social Anxiety Disorder. Psychoneuroendocrinology, 63, 26-33, doi:http://doi.org/10.1016/j.psyneuen.2015.09.008.

Fanning, J. R., Keedy, S., Berman, M. E., Lee, R., & Coccaro, E. F. (2017). Neural Correlates of Aggressive Behavior in Real Time: a Review of fMRI Studies of Laboratory Reactive Aggression. Current Behavioral Neuroscience Reports, 4(2), 138-150, doi:10.1007/s40473-017-0115-8.

Fox, M. D., Snyder, A. Z., Vincent, J. L., & Raichle, M. E. (2007). Intrinsic Fluctuations within Cortical Systems Account for Intertrial Variability in Human Behavior. Neuron, 56(1), 171-184, doi:http://dx.doi.org/10.1016/j.neuron.2007.08.023.

Frühholz, S., Klaas, H. S., Patel, S., & Grandjean, D. (2015). Talking in Fury: The Cortico-Subcortical Network Underlying Angry Vocalizations. Cerebral Cortex, 25(9), 2752-2762, doi:10.1093/cercor/bhu074.

Fulwiler, C. E., King, J. A., & Zhang, N. (2012). Amygdala-Orbitofrontal Resting State Functional Connectivity is Associated with Trait Anger. Neuroreport, 23(10), 606-610, doi:10.1097/WNR.0b013e3283551cfc.

Giancola, P. R., & Parrott, D. J. (2008). Further evidence for the validity of the Taylor Aggression Paradigm. Aggressive Behavior, 34(2), 214-229, doi:10.1002/ab.20235.

Gospic, K., Mohlin, E., Fransson, P., Petrovic, P., Johannesson, M., & Ingvar, M. (2011). Limbic Justice— Amygdala Involvement in Immediate Rejection in the Ultimatum Game. PLoS Biol, 9(5), e1001054, doi:10.1371/journal.pbio. 1001054.

Granger, D. A., Kivlighan, K. T., Fortunato, C., Harmon, A. G., Hibel, L. C., Schwartz, E. B., et al. (2007). Integration of salivary biomarkers into developmental and behaviorally-oriented research: Problems and solutions for collecting specimens. Physiology & Behavior, 92(4), 583-590, doi:http://dx.doi.org/10.1016/j.physbeh.2007.05.004.

Grèzes, J., Adenis, M.-S., Pouga, L., & Armony, J. L. (2013). Self-relevance modulates brain responses to angry body expressions. Cortex, 49(8), 2210-2220, doi:http://dx.doi.org/10.1016/j.cortex.2012.08.025.

Hahn, A., Kranz, G. S., Sladky, R., Kaufmann, U., Ganger, S., Hummer, A., et al. (2016). Testosterone affects language areas of the adult human brain. Human brain mapping, 37(5), 1738-1748, doi:10.1002/hbm.23133.

Hahn, A. C., Fisher, C. I., Cobey, K. D., DeBruine, L. M., & Jones, B. C. (2016). A longitudinal analysis of women’s salivary testosterone and intrasexual competitiveness. Psychoneuroendocrinology, 64, 117-122, doi:https://doi.org/10.1016/j.psyneuen.2015.11.014.

Herpertz, S. C., Nagy, K., Ueltzhöffer, K., Schmitt, R., Mancke, F., Schmahl, C., et al. (2017). Brain Mechanisms Underlying Reactive Aggression in Borderline Personality Disorder—Sex Matters. Biological Psychiatry, doi:https://doi.org/10.1016/j.biopsych.2017.02.1175.

Ho, N.-F., Chong, J. S. X., Koh, H. L., Koukouna, E., Lee, T.-S., Fung, D., et al. (2015). Intrinsic Affective Network Is Impaired in Children with Attention-Deficit/Hyperactivity Disorder. PLoS ONE, 10(9), e0139018, doi:10.1371/journal.pone.0139018.

Hoptman, M. J., D'Angelo, D., Catalano, D., Mauro, C. J., Shehzad, Z. E., Kelly, A. M. C., et al. (2010). Amygdalofrontal Functional Disconnectivity and Aggression in Schizophrenia. Schizophrenia Bulletin, 36(5), 1020-1028, doi:10.1093/schbul/sbp012.

Hull, J. V., Jacokes, Z. J., Torgerson, C. M., Irimia, A., & Van Horn, J. D. (2017). Resting-State Functional Connectivity in Autism Spectrum Disorders: A Review. Frontiers in Psychiatry, 7, 205, doi:10.3389/fpsyt.2016.00205.

Kaufman, M. J., Janes, A. C., Hudson, J. I., Brennan, B. P., Kanayama, G., Kerrigan, A. R., et al. (2015). Brain and Cognition Abnormalities in Long-Term Anabolic-Androgenic Steroid Users. Drug and Alcohol Dependence, 152, 47-56, doi:10.1016/j.drugalcdep.2015.04.023.

Keevil, B., MacDonald, P., Macdowall, W., Lee, D., & Wu, F. (2013). Salivary testosterone measurement by liquid chromatography tandem mass spectrometry in adult males and females. Annals of Clinical Biochemistry: An international journal of biochemistry and laboratory medicine, 51(3), 368-378, doi:10.1177/0004563213506412.

Low, K. L., Ma, C., & Soma, K. K. (2017). Tyramide Signal Amplification Permits Immunohistochemical Analyses of Androgen Receptors in the Rat Prefrontal Cortex. Journal of Histochemistry & Cytochemistry, 65(5), 295-308, doi:doi:10.1369/0022155417694870.

Madsen, M. K., Mc Mahon, B., Andersen, S. B., Siebner, H. R., Knudsen, G. M., & Fisher, P. M. (2016). Threat-related amygdala functional connectivity is associated with 5-HTTLPR genotype and neuroticism. Social Cognitive and Affective Neuroscience, 11(1), 140-149, doi:10.1093/scan/nsv098.

Maron-Katz, A., Vaisvaser, S., Lin, T., Hendler, T., & Shamir, R. (2016). A large-scale perspective on stress-induced alterations in resting-state networks. Scientific Reports, 6, 21503, doi:10.1038/srep21503.

McCloskey, M. S., Phan, K. L., Angstadt, M., Fettich, K. C., Keedy, S., & Coccaro, E. F. (2016). Amygdala hyperactivation to angry faces in intermittent explosive disorder. Journal of Psychiatric Research, 79, 34-41, doi:http://dx.doi.org/10.1016/i.jpsychires.2016.04.006.

Mehta, P. H., & Beer, J. (2009). Neural Mechanisms of the Testosterone-Aggression Relation: The Role of Orbitofrontal Cortex. Journal of cognitive neuroscience, 22(10), 2357-2368, doi:10.1162/jocn.2009.21389.

Mennes, M., Kelly, C., Colcombe, S., Castellanos, F. X., & Milham, M. P. (2013). The Extrinsic and Intrinsic Functional Architectures of the Human Brain Are Not Equivalent. Cerebral Cortex, 23(1), 223-229, doi:10.1093/cercor/bhs010.

Mothes-Lasch, M., Miltner, W. H. R., & Straube, T. (2012). Processing of angry voices is modulated by visual load. NeuroImage, 63(1), 485-490, doi:http://dx.doi.org/10.1016/j.neuroimage.2012.07.005.

Motzkin, J. C., Newman, J. P., Kiehl, K. A., & Koenigs, M. (2011). Reduced Prefrontal Connectivity in Psychopathy. The Journal of Neuroscience, 31(48), 17348-17357, doi:10.1523/jneurosci.4215-11.2011.

New, A. S., Hazlett, E. A., Buchsbaum, M. S., Goodman, M., Mitelman, S. A., Newmark, R., et al. (2007). Amygdala-Prefrontal Disconnection in Borderline Personality Disorder. Neuropsychopharmacology, 32(7), 1629-1640.

Peters, S., Jolles, D. J., Duijvenvoorde, A. C. K. V., Crone, E. A., & Peper, J. S. (2015). The link between testosterone and amygdala–orbitofrontal cortex connectivity in adolescent alcohol use. Psychoneuroendocrinology, 53, 117-126, doi:http://dx.doi.org/10.1016/j.psyneuen.2015.01.004.

Peters, S., Peper, J. S., van Duijvenvoorde, A. C. K., Braams, B. R., & Crone, E. A. (2016). Amygdala–orbitofrontal connectivity predicts alcohol use two years later: a longitudinal neuroimaging study on alcohol use in adolescence. Developmental Science, 20(4), 1-11, doi:10.1111/desc.12448.

Pope, J. H. G., Wood, R. I., Rogol, A., Nyberg, F., Bowers, L., & Bhasin, S. (2014). Adverse Health Consequences of Performance-Enhancing Drugs: An Endocrine Society Scientific Statement. Endocrine Reviews, 35(3), 341-375, doi:10.1210/er.2013-1058.

Prada, P., Hasler, R., Baud, P., Bednarz, G., Ardu, S., Krejci, I., et al. (2014). Distinguishing borderline personality disorder from adult attention deficit/hyperactivity disorder: A clinical and dimensional perspective. Psychiatry Research, 217(1), 107-114, doi:http://dx.doi.org/10.1016/j.psychres.2014.03.006.

Qin, S., Young, C. B., Duan, X., Chen, T., Supekar, K., & Menon, V. (2014). Amygdala Subregional Structure and Intrinsic Functional Connectivity Predicts Individual Differences in Anxiety During Early Childhood. Biological Psychiatry, 75(11), 892-900, doi:http://dx.doi.org/10.1016/j.biopsych.2013.10.006.

Rausch, J., Gäbel, A., Nagy, K., Kleindienst, N., Herpertz, S. C., & Bertsch, K. (2015). Increased testosterone levels and cortisol awakening responses in patients with borderline personality disorder: Gender and trait aggressiveness matter. Psychoneuroendocrinology, 55, 116-127, doi:http://dx.doi.org/10.1016/j.psyneuen.2015.02.002.

Ritter, D., & Eslea, M. (2005). Hot Sauce, toy guns, and graffiti: A critical account of current laboratory aggression paradigms. Aggressive Behavior, 31(5), 407-419, doi:10.1002/ab.20066.

Roy, A. K., Shehzad, Z., Margulies, D. S., Kelly, A. M. C., Uddin, L. Q., Gotimer, K., et al. (2009). Functional connectivity of the human amygdala using resting state fMRI. NeuroImage, 45(2), 614-626, doi:http://dx.doi.org/10.1016/j.neuroimage.2008.11.030.

Sadaghiani, S., Poline, J.-B., Kleinschmidt, A., & D’Esposito, M. (2015). Ongoing dynamics in large-scale functional connectivity predict perception. Proceedings of the National Academy of Sciences of the United States of America, 112(27), 8463-8468, doi:10.1073/pnas.1420687112.

Savic, I., Frisen, L., Manzouri, A., Nordenstrom, A., & Hirschberg, A. L. (2017). Role of testosterone and Y chromosome genes for the masculinization of the human brain. Human brain mapping, doi:10.1002/hbm.23483.

Schepeijans, F., Eickhoff, S. B., Hömke, L., Mohlberg, H., Hermann, K., Amunts, K., et al. (2008). Probabilistic Maps, Morphometry, and Variability of Cytoarchitectonic Areas in the Human Superior Parietal Cortex. Cerebral Cortex, 18(9), 2141-2157, doi:10.1093/cercor/bhm241.

Sohn, W. S., Yoo, K., Lee, Y.-B., Seo, S. W., Na, D. L., & Jeong, Y. (2015). Influence of ROI selection on resting state functional connectivity: an individualized approach for resting state fMRI analysis. Frontiers in Neuroscience, 9, 280, doi:10.3389/fnins.2015.00280.

Spielberg, J. M., Forbes, E. E., Ladouceur, C. D., Worthman, C. M., Olino, T. M., Ryan, N. D., et al. (2015). Pubertal testosterone influences threat-related amygdala–orbitofrontal cortex coupling. Social Cognitive and Affective Neuroscience, 10(3), 408-415, doi:10.1093/scan/nsu062.

Sussman, T. J., Weinberg, A., Szekely, A., Hajcak, G., & Mohanty, A. (2016). Here Comes Trouble: Prestimulus Brain Activity Predicts Enhanced Perception of Threat. Cerebral Cortex, doi:10.1093/cercor/bhw104.

Terburg, D., Syal, S., Rosenberger, L. A., Heany, S. J., Stein, D. J., & Honk, J. v. (2016). Testosterone abolishes implicit subordination in social anxiety. Psychoneuroendocrinology, 72, 205-211, doi:http://doi.org/10.1016/j.psyneuen.2016.07.203.

Terburg, D., & van Honk, J. (2013). Approach–Avoidance versus Dominance–Submissiveness: A Multilevel Neural Framework on How Testosterone Promotes Social Status. Emotion Review, 5(3), 296-302, doi:10.1177/1754073913477510.

Tremblay, P. F., Graham, K., & Wells, S. (2008). Severity of physical aggression reported by university students: A test of the interaction between trait aggression and alcohol consumption. Personality and individual Differences, 45(1), 3-9, doi:http://dx.doi.org/10.1016/j.paid.2008.02.008.

Van den Stock, J., Hortensius, R., Sinke, C., Goebel, R., & de Gelder, B. (2015). Personality traits predict brain activation and connectivity when witnessing a violent conflict. [Article]. Scientific Reports, 5, 13779, doi:10.1038/srep13779.

van Wingen, G., Mattern, C., Verkes, R. J., Buitelaar, J., & Fernández, G. (2010). Testosterone reduces amygdala–orbitofrontal cortex coupling. Psychoneuroendocrinology, 35(1), 105-113, doi:10.1016/j.psyneuen.2009.09.007.

Vigil-Colet, A., Ruiz-Pamies, M., Anguiano-Carrasco, C., & Lorenzo-Seva, U. (2012). The impact of social desirability on psychometric measures of aggression. Psicothema, 24(2), 310-315.

Wahlund, K., & Kristiansson, M. (2009). Aggression, psychopathy and brain imaging — Review and future recommendations. international Journal of Law and Psychiatry, 32(4), 266-271, doi:http://dx.doi.org/10.1016/j.ijlp.2009.04.007.

Wang, J., Yang, Y., Fan, L., Xu, J., Li, C., Liu, Y., et al. (2015). Convergent functional architecture of the superior parietal lobule unraveled with multimodal neuroimaging approaches. Human brain mapping, 36(1), 238-257, doi:10.1002/hbm.22626.

Welker, K. M., Norman, R. E., Goetz, S., Moreau, B. J. P., Kitayama, S., & Carré, J. M. (2017). Preliminary evidence that testosterone's association with aggression depends on self-construal. Hormones and Behavior, 92, 117-127, doi:http://dx.doi.org/10.1016/j.yhbeh.2016.10.014.

Westlye, L. T., Kaufmann, T., Alnæs, D., Hullstein, I. R., & Bjernebekk, A. (2017). Brain connectivity aberrations in anabolic-androgenic steroid users. NeuroImage: Clinical, 13, 62-69, doi:http://dx.doi.org/10.1016/j.nicl.2016.11.014.

White, S. F., VanTieghem, M., Brislin, S. J., Sypher, I., Sinclair, S., Pine, D. S., et al. (2015). Neural Correlates of the Propensity for Retaliatory Behavior in Youths With Disruptive Behavior Disorders. American Journal of Psychiatry, 173(3), 282-290, doi:10.1176/appi.ajp.2015.15020250.

Whitfield-Gabrieli, S., & Nieto-Castanon, A. (2012). Conn: a functional connectivity toolbox for correlated and anticorrelated brain networks. Brain Connectivity, 2, 125-141, doi:10.1089/brain.2012.0073.

Yoder, K. J., Porges, E. C., & Decety, J. (2015). Amygdala subnuclei connectivity in response to violence reveals unique influences of individual differences in psychopathic traits in a nonforensic sample. Human brain mapping, 36(4), 1417-1428, doi:10.1002/hbm.22712.

Zhou, M., Hu, X., Lu, L., Zhang, L., Chen, L., Gong, Q., et al. (2017). Intrinsic cerebral activity at resting state in adults with major depressive disorder: A meta-analysis. Progress in Neuro-Psychopharmacology and Biological Psychiatry, 75, 157-164, doi:http://dx.doi.org/10.1016/j.pnpbp.2017.02.001.

Zuo, X.-N., Di Martino, A., Kelly, C., Shehzad, Z. E., Gee, D. G., Klein, D. F., et al. (2010). The Oscillating Brain: Complex and Reliable. NeuroImage, 49(2), 1432-1445, doi:10.1016/j.neuroimage.2009.09.037.

